# Translational Pharmacokinetics and Pharmacodynamics of a Cationic mRNA–Lipid Nanoparticle from Mice to Non Human Primates

**DOI:** 10.64898/2026.09.23.749423

**Authors:** Charlotte Maeve Dunne, Katrin Radloff, Leonidas Gkionis, Nourhan Kahwaji, Natascha Hartl, Oliver Keil, Angela Fischer, Gerrit Maass, Ansgar Santel, Daniel Tondera, Jörg Kaufmann

## Abstract

Cationic lipid nanoparticles have demonstrated unique potential for extrahepatic mRNA delivery, particularly enabling selective targeting of the pulmonary endothelium. However, their translational development has been hampered by reports of infusion-related immune reactions and innate immune system activation, most notably transient complement activation. Here, we present a case study illustrating the discovery and translational advancement of a selected cationic LNP into non-human primates (NHPs) for initial pharmacokinetic assessment and evaluation of potential immunostimulatory side effects. We show surface charge dependent organ-selective expression of reporter mRNAs from different LNPs *in vivo*. An mRNA encoding the Tie2 agonist COMP-Angl, was formulated with LNP002, and respective pharmacokinetic and pharmacodynamic readouts were analyzed in two independent non-human primate studies. Notably, dose-dependent transient complement activation could be abrogated by extending the infusion time. Finally, we identified the blood-borne pharmacodynamic biomarker PDGFB for LNP002/mRNA-76 treatment reflecting activated Tie2-signalling in healthy pulmonary endothelium *in vivo* supported by single cell sequencing and cluster-alignment of downstream effector genes with the same spatial profile as the delivered mRNA.

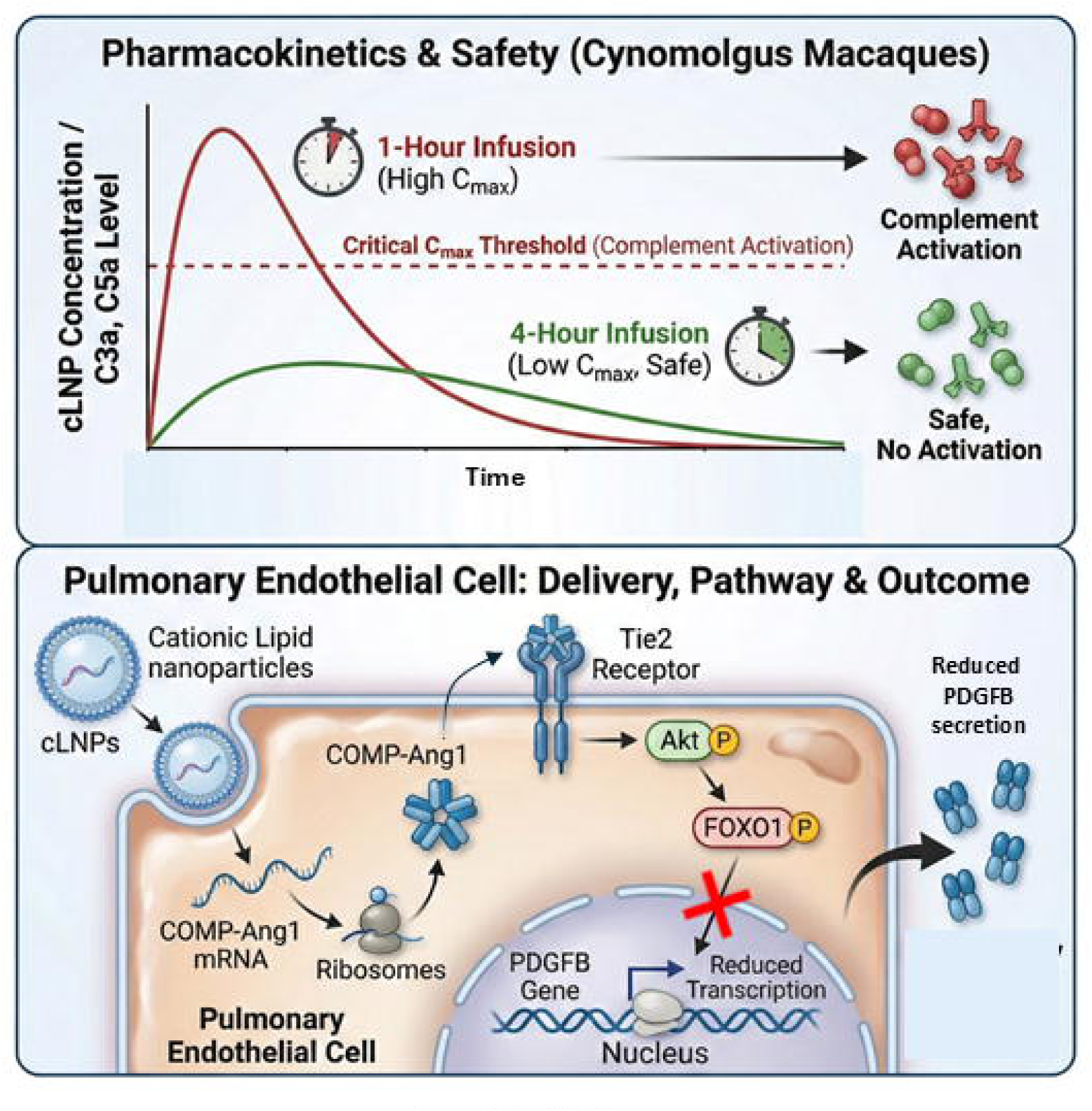

## Introduction

The clinical translation of gene therapy mediated by DNA- or RNA-based modalities has historically been restrained by the lack of effective delivery systems. Viral vectors are limited by the size of the encapsulated DNA or RNA, and by concerns regarding immunogenicity and safety due to respective random integration into the target-genome. Applications of non-viral delivery systems have long since been hampered by the lack of potency and safety concerns. However, the technology for non-viral lipid nanoparticles has advanced dramatically within the last decade as shown by clinically approved siRNA therapeutics and numerous other candidates in clinical trials reviewed elsewhere.^1–3^ The success of the mRNA-based COVID-19 vaccines has also highlighted the transforming character of mRNA-LNP technologies and is currently being expanded into treatment for genetic diseases, immunotherapies for cancer and for other indications.^4–6^ To bring the potential of non-viral delivery systems to fruition, further LNP technology developments will be required in order to improve potency and therapeutic index, to expand tissue and cell selectivity and additionally to increase safety.

To date, all 5 clinically approved RNA-LNP formulations^4,5,7,8^ rely on ionizable lipids and exhibit a pronounced tropism to the liver following intravenous administration, this being largely mediated by apolipoprotein E (ApoE)–dependent uptake in hepatocytes.^9,10^ While this property has facilitated the development of liver-directed therapies, it has simultaneously restricted broader application of RNA therapeutics to extrahepatic targets. Cationic lipid nanoparticles (cLNPs), typified by permanently positively charged cationic lipids i.e. lipids substantially protonated throughout the physiologically relevant pH-range, can be generated by using a molar excess of positive lipid charges over the negative nucleic acid charges. In doing so, these cLNPs acquire a slightly overall positive particle surface charge (positive zeta-potential), thus becoming a distinct class of delivery systems exhibiting fundamentally different biodistribution characteristics when compared to classical neutral LNPs for hepatic delivery. Especially particles with positive surface charge, including cLNPs, have repeatedly been shown to preferentially target the pulmonary vasculature following systemic administration.^11–15^ We have recently reported preclinical data for a cLNP, “LNP002” containing a permanently positive charged lipid, (L-Arginyl)-L-2,3-diamino propionic acid-N-palmityl-N-oleyl-amide, to encapsulate mRNA-76 encoding COMP-Ang-1, a Tie-2 agonist preventing pulmonary vascular leakage. The cationic delivery system, LNP002 in this study was very effective in transporting mRNAs selectively and almost exclusively into the lung capillary endothelial cells (ECs). Systemic bolus tail-vein administration led to robust mRNA mediated expression of luciferase reporter or COMP-Ang-1 in mice. However, despite their favorable and exclusive lung targeting profile, cLNPs have a poor reputation for translational applications. This is in part due to the widespread and false assumption that permanently charged cationic lipids within LNP formulations, such as those employed in early studies with DNA lipoplexes, should be avoided due to safety issues.^16^ Indeed several *in vivo* preclinical studies with DNA plasmid or antisense payloads have described toxicities induced by systemic administration in mice, including activation of the complement system as well as acute inflammatory responses.^17–20^ These findings have contributed to the perception that cationic lipid chemistry, employing permanently positively charged headgroups possibly inherently limits systemic tolerability. Notably, however, most of these observations originate from experimental settings characterized by bolus-injections in rodents, or by short- duration intravenous infusions in non-human primates (NHPs), conditions that generate very high peak plasma concentrations (C_max_), and thus may not adequately reflect clinically relevant dosing paradigms. Infusion-related reactions to nanoparticle-based therapeutics are well acknowledged across multiple drug classes, including liposomal formulations and biologics, and are frequently driven by peak exposure rather than by cumulative dose.^21^ Interestingly, the only currently approved neutral LNP (nLNP) based nucleic acid systemic therapy, Patisiran/Onpattro siRNA (Alnylam) is based on the ionizable lipid MC-3, does indeed require an extensive premedication to mitigate adverse immune stimulation in patients. With regard to safety, the drug is injected in combination with paracetamol, antihistamines, corticosteroids, and ranitidine to prevent infusion-related adverse reactions.^8,22^ Whether the immune activation observed with cationic and/or neutral LNPs reflects intrinsic and specifically lipid-compound-mediated toxicity, or rather represents a C_max_-dependent “foreign surface” phenomenon, remains insufficiently explored.

In this study, we aimed to characterize the pharmacokinetics, pharmacodynamics, and preliminary tolerability of the cationic mRNA-COMP-Ang1-formulation “LNP002/mRNA-76” as a foundation for future preclinical translation and subsequent GLP toxicity testing.

## Results

### Charge-dependent tropism confirms suitability of cationic LNPs for pulmonary mRNA delivery

Initial experiments compared the transfection efficiency and biodistribution of cationic and neutral LNP formulations encapsulating reporter mRNA. *In vitro* transfection of HeLa cells demonstrated robust, concentration-dependent reporter expression mediated by cLNPs, whereas most nLNPs showed markedly reduced activity (**Figure 1A**). These data corroborate the observation that most commercial transfection reagents for *in vitro* studies which employ adherent growing eukaryotic cells, are based on positively charged components such as calcium phosphate and in subsequent years, cationic lipids. In our experiments, no substantial differences in overall cell morphology were observed among these classes of LNPs i*n vitro* as seen in phase contrast images (**Figure 1A**). Systemic administration in mice resulted in pronounced lung-restricted expression following delivery with cLNPs, while nLNPs predominantly mediated hepatic expression (**Figure 1B**). The overall cLNP-mediated luciferase reporter expression level in the lung tissue was around 5-10 times higher than the corresponding liver expression when normalized for tissue weight. This observed difference in expression in whole tissue lysates likely results from the higher potency of the ApoE dependent uptake of nLNPs by liver cells as well as from the predominance of hepatocytes and macrophages as the main cell types in the liver. In contrast, uptake of cLNPs in the lung is selective to endothelial cells of the lung vasculature (General /alveolar capillaries) (see **Figure 5C**). These findings confirm the charge-dependent tropism of cationic LNPs, as previously described,^23^ and reinforce their suitability as delivery vehicles for pulmonary endothelial-targeted mRNA therapeutics.

**Figure 1.**
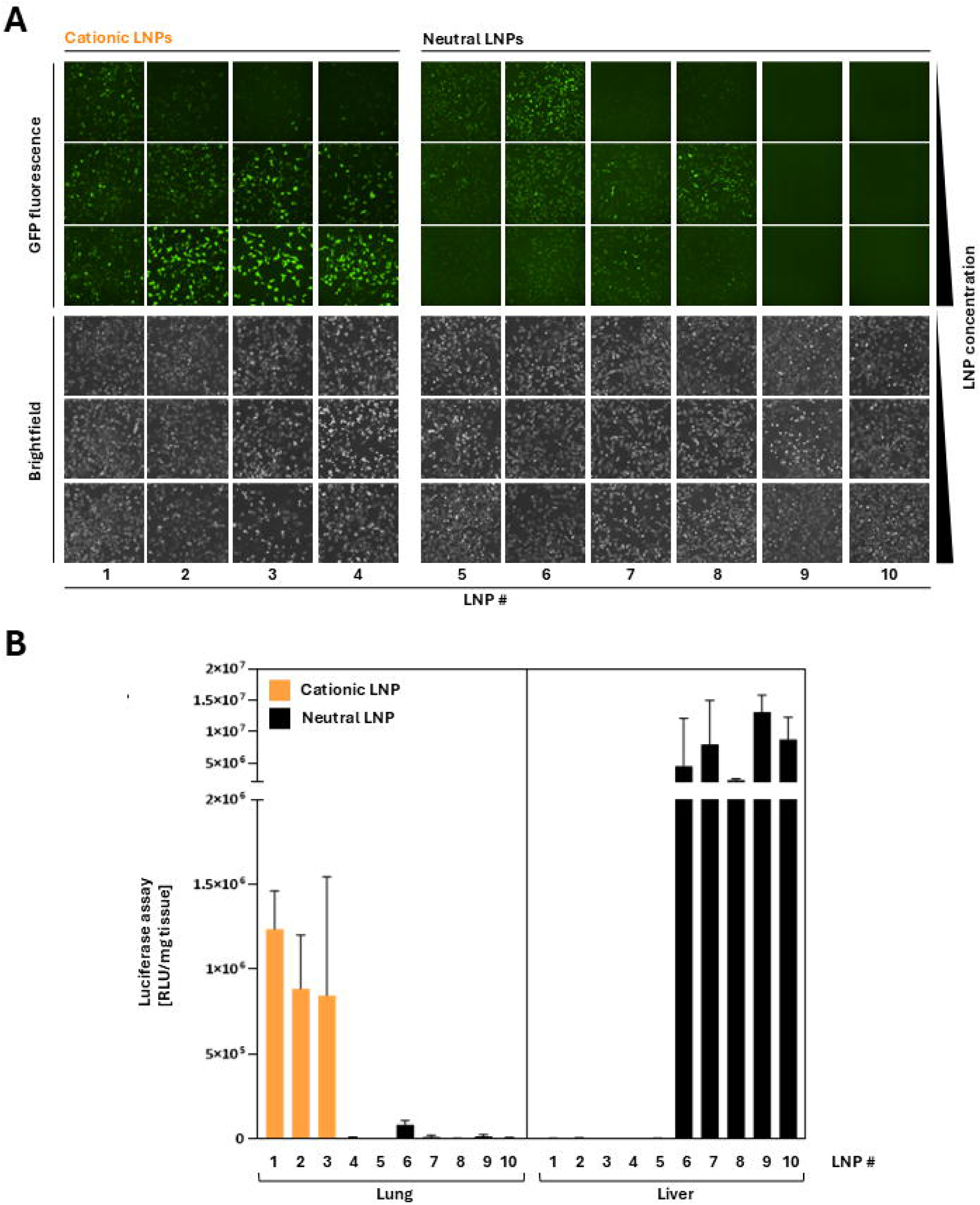
Surface charge-dependent tropism of lipid nanoparticle (LNPs) mediated mRNA delivery in vitro and in vivo. **(A)** Comparison of mRNA expression mediated by cationic LNPs with classical neutral LNPs in vitro. Representative fluorescence microscopy images of HeLa cells following transfection with GFP-encoding mRNA formulated in either cationic (LNP 1-4) or neutral lipid nanoparticles (LNP 5-10). Increasing LNP concentrations of (0.25, 0.5 and 1 μg/mL) were used for cationic LNPs whereas (2, 10, 25 μg/mL) were used for neutral LNPs which typically require a higher concentration in vitro compared to cationic LNPs. GFP fluorescence indicates transgene expression, while corresponding brightfield images are shown to control for cell morphology and viability. **(B)** In vivo biodistribution of cationic (orange) and neutral (black) LNPs in the lung and liver 4 hr post bolus tail vein intravenous administration of 1.5 mg/kg luciferase mRNA encapsulated in LNPs in C57BI/6NRj male mice (n=3). Tissues were harvested, homogenized and analyzed by luciferase assay.

### Upscaling of LNP002 formulation accompanied by physicochemical analysis supports translation into *in vivo* studies

We generated several batches of LNP002/mRNA-76 using a microfluidic mixer (NanoAssemblr, Precision Nanosystem) as described previously.^23^ Different batches of the LNP002/mRNA-76 formulation were tested for particle size, amounting to very low variation between 80 and 90 nm. Dynamic light scattering analysis demonstrated consistent particle-size distributions across independent formulation batches, indicating excellent manufacturing reproducibility (**Figure 2A**). Particle size remained stable following freeze–thaw cycles (**Figure 2B**) and after dilution in saline under relevant conditions (to 30, 60 and 90 µg/ml mRNA) for intravenous infusion in NHPs (**Figure 2C-E**). Collectively, these data demonstrate robust physicochemical stability of the cationic LNP002/mRNA-76 across manufacturing, storage, and dilution conditions required for translational in vivo applications.

**Figure 2.**
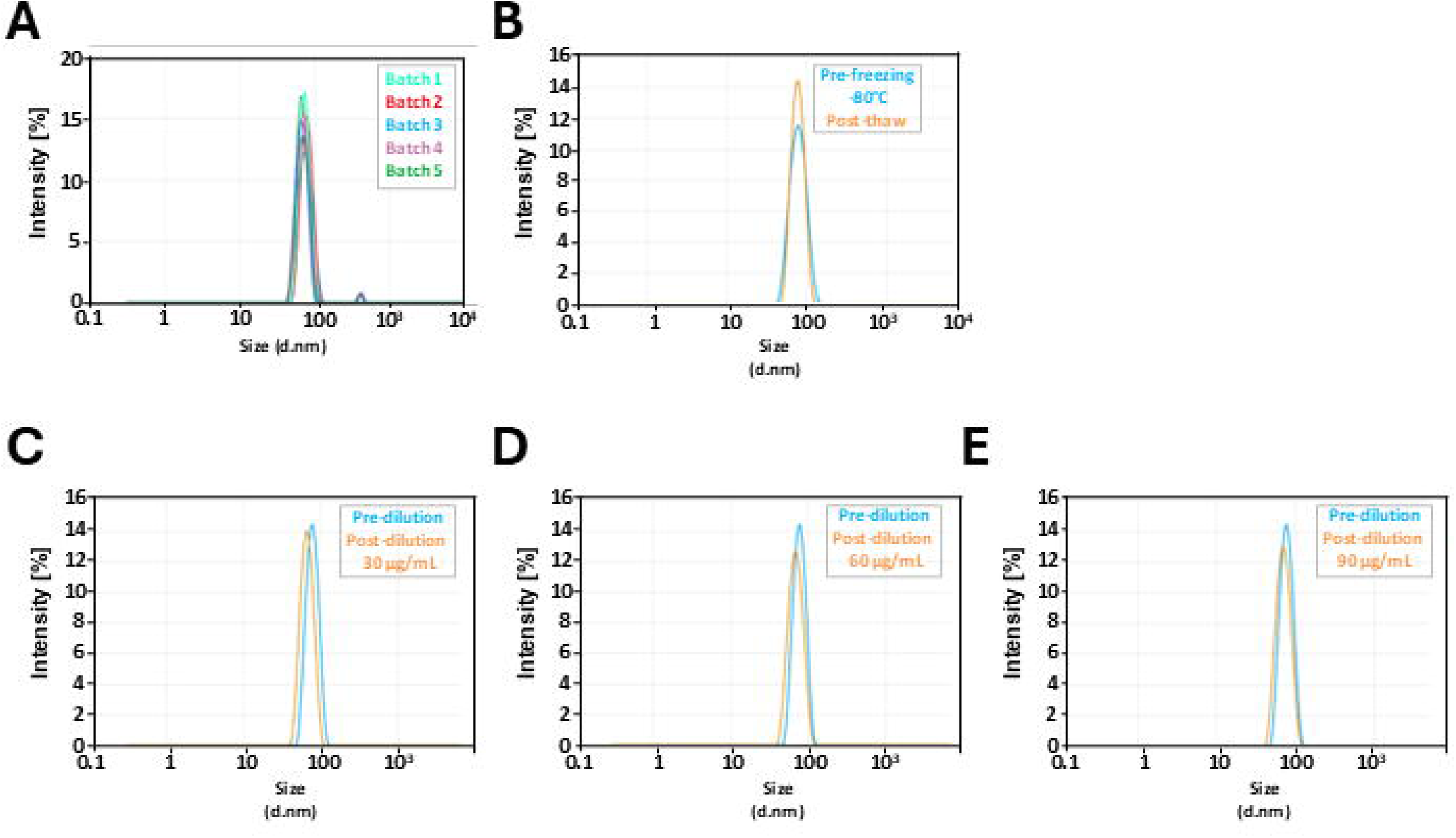
Physicochemical characterization of the selected cationic LNP002. Dynamic light scattering (DLS) was used to assess the hydrodynamic diameter and size distribution of the selected cationic lipid nanoparticle formulation LNP002 under different manufacturing and handling conditions. **(A)** Batch-to-batch consistency of LNP002 particle size obtained from five independent formulation runs. **(B)** Stability of LNP002 following storage at −80 °C, comparing particle size distributions of freshly formulated LNPs prior to freezing with those measured after thawing. **(C-E)** Particle size distribution of LNP002 before (200 µg/mL) and after dilution in saline to 30, 60 and 90 µg/mL, reflecting formulation stability under conditions relevant for subsequent in vivo administration in non-human primates (NHP).

### Pharmacokinetic (PK) NHP studies comparing 1-hour and 4-hour infusions to assess C_max_- driven immune responses

As a preclinical development candidate, LNP002/mRNA-76 is designed for intravenous (i.v.) administration in hospital ICUs to prevent lung edema formation in acute respiratory distress syndrome (ARDS) patients via activation of Tie-2, which in turn counteracts inflammatory vascular leakage.^23^ Hence, we evaluated the suitability of LNP002 for future clinical use and set out to investigate the impact of infusion kinetics on immunological tolerability. Adverse infusion reactions are a well-known feature of particle-based drugs including LNPs and need to be well-considered early in the drug development process. For this reason, we analyzed the cytokine response and complement activation in mice after bolus administration of LNP002/mRNA^Luc^. Doses known to be well tolerated in mice, even when administered repeatedly, were chosen and an expected strong cytokine and complement response were observed (**see Supplemental Figure 1**). Surprisingly, cytokine responses and complement activation were both observed to be transient and had no significant effect on murine body weight or on respective general wellbeing, this indicated good tolerability despite the high C_max_ owing to bolus administration. However, as a cautionary note, the apparent tolerability observed in mice likely reflects the fact that rodents, especially mice, are poor models for predicting immune toxicity or complement.^24^ For a more relevant assessment, two pilot PK studies in non-human primates (NHP) were conducted using cationic LNP002/mRNA-76 formulations, applying single-drug administrations and different durations of infusion. Naïve as well as pre-treated NHPs were enrolled in these two non-GLP- conform studies. Animals received i.v. administration either as a short 1-h infusion, or as a prolonged 4-h infusion at mounting doses of 0.3, 0.6, and 0.9 mg/kg mRNA, with serial blood sampling performed before and after dosing (**Figure 3 A and B**). This design allowed a direct comparison of mRNA PKs, as well as assessment of blood mRNA levels, and immune activation profiles across different infusion durations, while maintaining total administered doses. The PK evaluation showed dose-dependent exposure across all doses for both the 1- and 4-h infusion, with no notable differences in PK profiles between the tested dose concentrations (**Figure 3 C and D**). The C_max_ values for all treatments occurred immediately at the end of infusion, regardless of dose or infusion duration, indicating rapid mRNA clearance of non-delivered LNPs from serum. This suggests both efficient cellular uptake and additional non-specific absorption and degradation processes. Our study design enabled a direct comparison of C_max_ across different infusion durations, and the effect of reduced a C_max_ on subsequent immunological responses supporting the assessment of infusion time as a potential mitigation strategy for infusion-related reactions.

**Figure 3.**
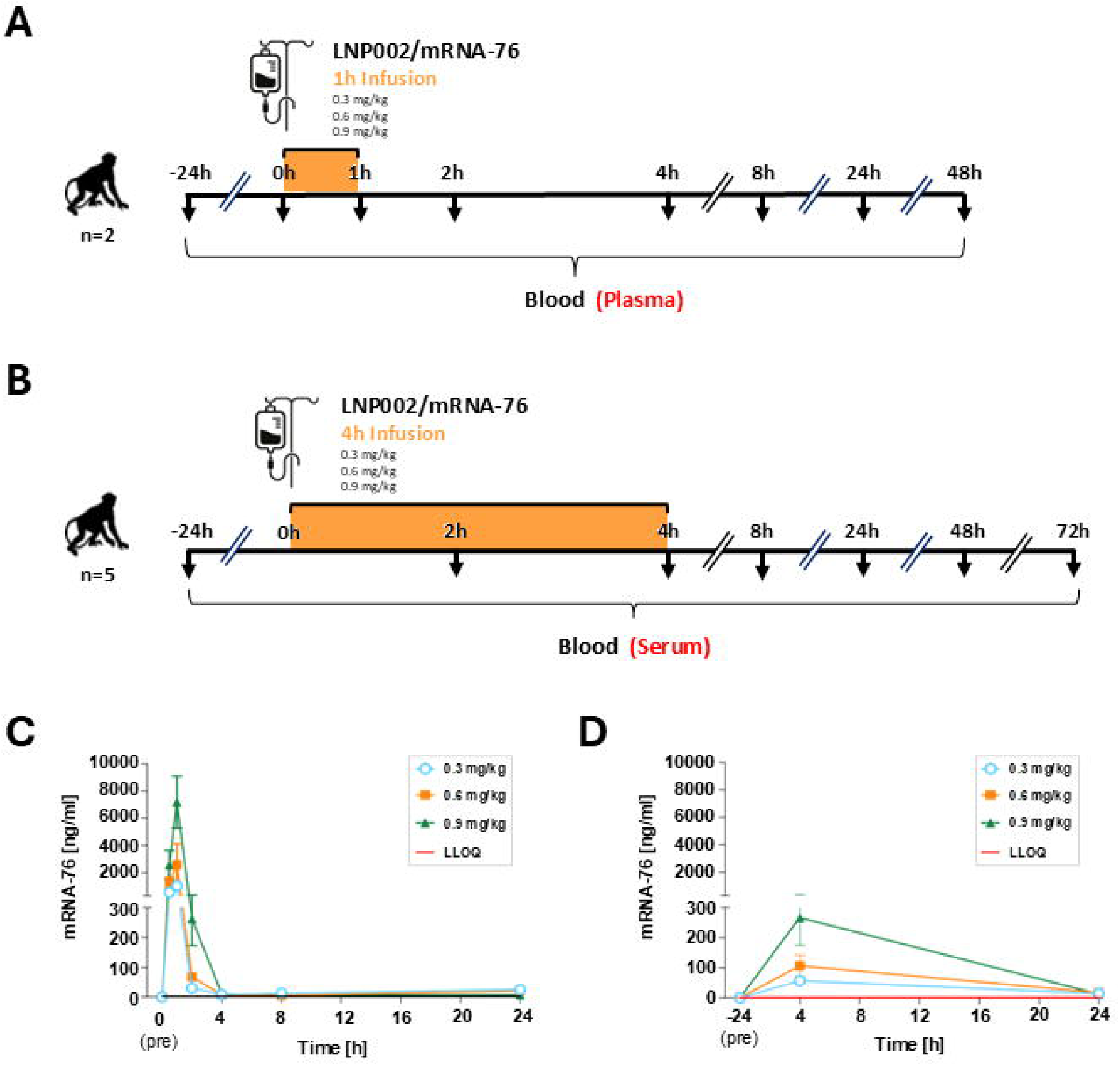
Pharmacokinetic studies with LNP002/mRNA-76 in non-human primates (NHP) to assess mRNA serum levels over time **(A)** Schematic overview of the treatment and sampling schedules of non-naïve male NHPs (n = 2) received LNP002/mRNA-76 via intravenous infusion over 1 h whereas in **(B**) Non-naïve male NHPs (n = 5) received LNP002/mRNA-76 via prolonged intravenous infusion over 4 h. Infusion time dependent plasma mRNA levels are shown in **(C)** for 1 h infusion and **(D)** for 4 h infusion. For both studies the C_max_ is reached immediately at the end of the infusion indicating a very rapid clearance or uptake of the mRNA.

### Mitigation of C_max_-Driven Infusion Reactions Enables safe systemic administration of cationic lipid nanoparticles in NHPs

Having confirmed drug exposure in all animals, characterized by a rapid peak in plasma levels of encapsulated mRNA with clear dose dependence, we next investigated immune activation in these animals. Cytokine responses were observed to be minimal and insignificant for both the 1- h and the 4-h infusions (**Figure 4A and 4B**). There was no clear dose-dependence of MCP-1 and a weak but transient IL-6 induction in only the high dose groups (0.9 mg/kg) in the 1-h perfusion study with a lower IL-6 induction in the 4-h perfusion study. No consistent dose or time dependent induction of pro-inflammatory cytokines (including, IL-8, TNFα, IL-4 or IL-5) were observed. No signal was detected for IL-10, IL-1b, IL-2, IL-17 nor for IFN-γ in both studies. Following an increase in the infusion time to 4 h, cytokine profiles remained minimal and comparable to baseline (**Figure 4B**), further confirming the immunological tolerability associated with cationic lipid nanoparticles.

**Figure 4.**
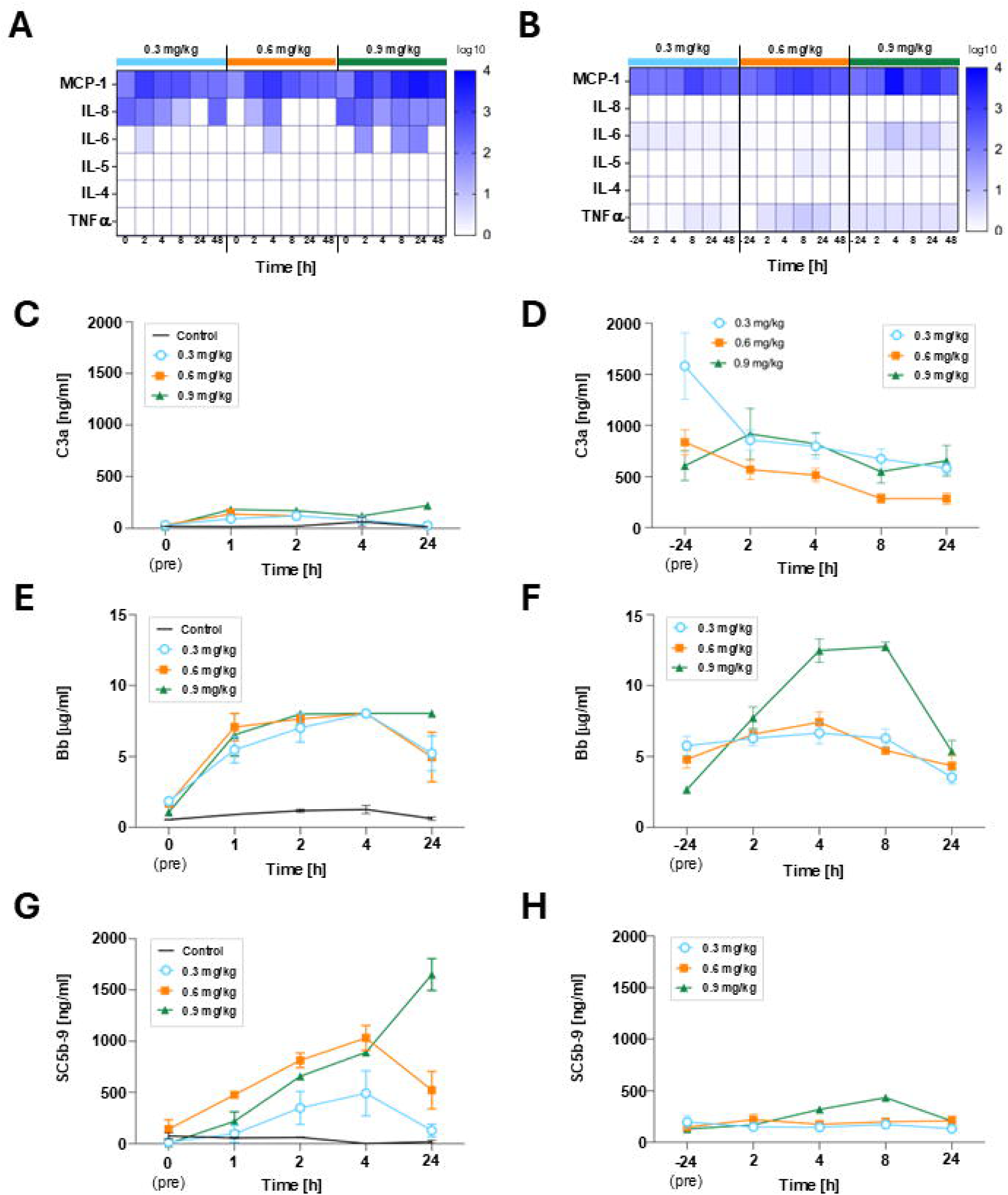
Cytokine response and complement activation after infusion of a single LNP002/mRNA-76 dose in non-human primates Cytokine response and complement activation were assessed in two NHP studies comparing 1 h versus a prolonged 4 h intravenous infusion of cationic LNP formulations at increasing dose levels (0.3, 0.6, and 0.9 mg/kg). Selected cytokine and chemokines responses presented as heat maps for **(A)** 1h infusion time **(B)** 4h infusion time. Complement activation was investigated by analyzing C3a levels **(C), (D)** reflecting an upstream complement activation, Bb fragment **(E), (F)** as an indicator of alternative pathway activation and Soluble C5b-9 (sC5b-9) **(G), (H)** as a marker of terminal complement activation.

**Figure 5.**
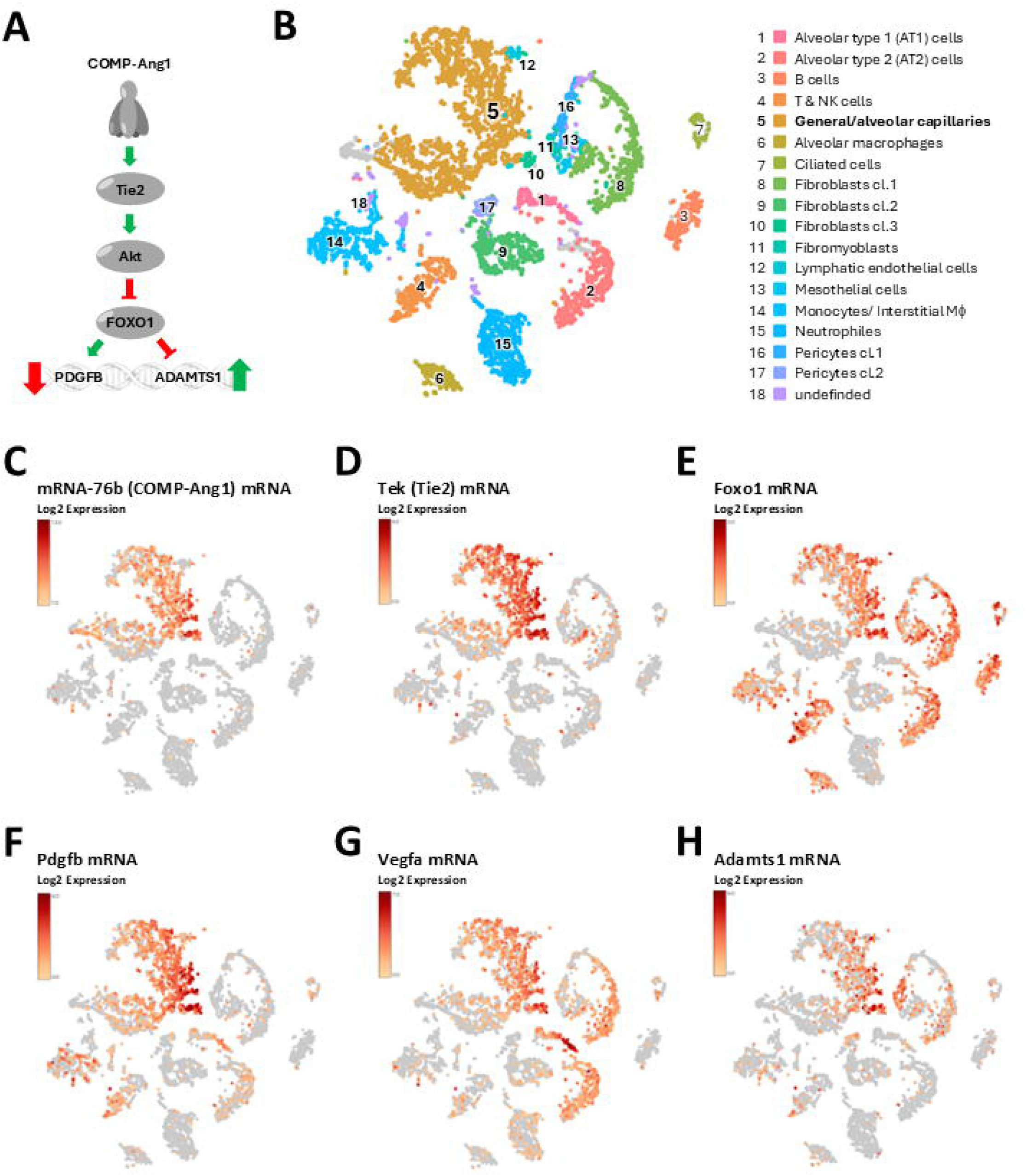
Identification of potential Tie2 signaling downstream targets by single-cell RNA sequencing analysis and colocalized expression with Tie2 receptor and mRNA-76b **(A)** Schematic signal transduction cascade of the Tie2 pathway predicting potential upregulated or downregulated effector mRNAs and their encoded proteins. **(B)** Loupe- based t-SNE clustering of murine lung cell populations derived from single-cell RNA sequencing analysis, identifying 18 distinct cell-type clusters as indicated. **(C)** t-SNE feature plots showing the LNP002 mediated uptake and distribution of the synthetic mRNA-76 encoding COMP-Ang1in lung tissue of mice (single bolus administration). Cells annotated to Cluster 5 (general/alveolar capillaries) show the highest mRNA-76 uptake confirming the lung endothelium as the preferred target structure. (**D), (E) & (F)** t-SNE feature plots depicting expression levels of endogenous mRNAs which are colocalized lung endothelial cells derived from capillaries (cluster 5). The endogenous mRNA expression of COMP-Ang1 receptor Tie2 **(D)**, VEGF-A **(G)** and the previously reported downstream Tie2 pathway targets; PDGFB **(F)** and Adamts1 **(H)** are all annotated to cell-type cluster 5. Expression intensity is shown as log2-scaled signal (red).

### Short infusion time of a high dose of LNP002/mRNA-76 can induce complement activation

Complement induction has been reported for both neutral^20,25^ and cationic LNPs.^17,19,20^ We therefore investigated the blood from both studies by analyzing the changes of the general complement activation marker C3a, of the alternative pathway marker Bb, and of the terminal pathway marker sC5b-9 (**Figure 4**). Following the 1-h infusion, an activation of the complement system was observed, as demonstrated by transient increases in levels of sC5b-9, Bb, and C3a for the 0.3 mg/kg and 0.6 mg/kg doses (**Figure 4C, E & G**). In contrast, the complement activation of all three markers for the 0.9 mg/kg dose of LNP002/mRNA-76 applied in a 1-h infusion setting was not transient. Notably, the highest level of the terminal complement pathway marker sC5b-9 was observed at the latest analyzed timepoint, 24 h after infusion-start. This pronounced elevation of the terminal complement cascade marker, sC5b-9 (**Figure 4G**), indicates full activation of the complement cascade and a corresponding risk of tissue damage.

### Prolonged infusion time markedly reduces complement activation despite comparable exposure

Extension of the infusion duration to 4 h resulted in altered PK, characterized by reduced peak plasma concentrations, while maintaining overall exposure (**Figure 3D**). Under these conditions, complement activation was markedly attenuated or completely absent across all tested dose levels (**Figure 4D, F & H**). Only the high dose of LNP002/mRNA-76 induced a transient and reversible increase in levels of Bb and sC5b-9, peaking at 8 hours and returning to baseline by 24 h. Complement activation correlated with administered dose and plasma concentration profile, consistent with a C_max_-driven mechanism. Hence, this reduction or prevention of complement activation across all dose levels, despite comparable overall exposure can be mitigated by prolonging the duration of infusion.

### Identification of COMP-Ang1/Tie2-Dependent Secreted PD Biomarker Candidates

In the initial PK/PD analysis, we aimed at evaluating pharmacodynamic endpoints. A robust COMP-Ang-1 expression in murine lung tissue lysates was previously demonstrated^23^ but a secreted and ideally lung-borne PD-marker for endothelial Tie2-activation, detectible in blood is yet to be identified. Unfortunately, a direct detection of secreted COMP-Ang-1 monomer in blood or serum by immunoblot or ELISA following LNP002/mRNA-76 treatment has proven challenging despite confirmed expression in lung tissue (data not shown). These detection challenges may be attributable to the extremely short half-life of recombinant COMP-Ang-1 in blood,^26^ or to the retention of COMP-Ang-1 in lung tissue with limited systemic release. For native Ang-1, high affinity binding to the heparan-sulfate, a key glycocalyx component, along with strong extracellular matrix interactions have been described, and could explain this retention.^27,28^ To identify a new secreted PD marker we examined blood-based markers downstream of COMP-Ang-1 mediated Tie2 activation under non-inflammatory conditions. We hypothesized that the lung-vascular specific activation of this pathway might alter the expression and secretion of specific downstream proteins, enabling their detection in blood (see proposed Tie2 signaling cascade is presented in **Figure 5A**), indicating genes whose expression is either upregulated or downregulated in response to Tie2 activation. COMP-Ang-1 was initially proposed to most likely act in an autocrine manner in endothelial cells transfected by LNP002/mRNA-76 but could also mediate Tie2 activation in a paracrine manner. We re- analyzed rodent scRNA-seq data of lung tissue from mice treated with LNP002/mRNA-76 to assess the spatial relevance of these downstream Tie 2 receptor targets *in vivo*.

t-SNE clustering revealed 18 distinct cell populations (**Figure 5B**). Single-cell transcriptomic analysis confirmed the precise co-localization of mRNA-76 (**Figure 5C**) and Tie2 mRNA (**Figure 5D)** within the alveolar capillary endothelial cluster, enabling potential autocrine COMP-Ang1 engagement of Tie2 signaling. LNP002/mRNA-76 mediated Tie2-activation was previously shown in lung tissue of mice.^23^ Expression of mRNAs encoding the transcription factor Foxo1 and secreted proteins PDGFB, VEGFA and ADAMTS1 was also localized to the same mRNA-76/Tie2 cluster (5), representing blood capillaries of the lung (**Figure 5E-H**). Notably, secreted PDGFB and ADAMTS1 exhibited highly spatially restricted expression-patterns within this mRNA-76/Tie2 cluster, positioning both as strong candidates for PD marker of COMP-Ang1-mediated Tie2-activation. Moreover, both genes, PDGFB and ADAMTS1, have been previously reported to show either downregulated or upregulated mRNA expression in endothelial cells, or in tissue derived from venous malformation patients with hyperactivating Tie2 mutation^29,30^ making them suitable candidates for further studies.

### Identification of PDGFB blood levels as a secondary PD marker for LNP002/mRNA-76 induced Tie2 activation in mice and NHPs

We identified ADAMTS1 and PDGFB by scRNA-seq to have endothelial-specific expression in lung tissue. However, since their expression is not restricted to the pulmonary vascular bed, systemic secretion from other tissues could potentially mask lung-specific changes. To rigorously test whether changes in protein levels can be induced by Tie2-activation in the lung, we treated mice with the highest tolerable dose of LNP002/mRNA-76 using various repeated dosing schedules (**Figure 6A**). No effect on body weight was detected, even in the group of mice receiving four bolus treatments (doses 0.75 mg/kg) on four consecutive days, again indicating tolerability of LNP002/mRNA-76 (**Figure 6B**). To validate PDGFB and ADAMTS1 as biomarkers for Tie2-activation in lungs, we performed ELISAs with serum derived from blood 24 h after the last respective treatment. Unfortunately, we were not able to generate reliable data for ADAMTS1 with different commercial ELISA systems nor with immunoblots employing different commercial antibodies. Conversely, ELISAs for serum PDFGB revealed a dose-dependent reduction in PDGFB protein levels in all groups, with the exception of the single-treatment group. The most robust reduction was observed with 4 consecutive daily treatments (**Figure 6C**), reinforcing a mechanistic explanation in which COMP-Ang1 suppresses PDGFB secretion. Unchanged serum VEGFA protein levels served as a negative control, as VEGF- expression and -secretion are not associated with Tie2-signaling **(Figure 6D)**. As mentioned above, we currently do not have analytical tools (e.g. ELISA, antibody) for reliable detection of COMP-Ang-1 in blood (serum or plasma), while a robust dose dependent COMP-Ang-1 expression can be demonstrated in murine lung lysates by immunoblot.^23^ We obtained lung tissue specimen from one (0.9 mg/kg) treated and non-treated NHP for subsequent immunoblot analysis with whole tissue protein lysates. Similar to lung lysates from the mouse studies, we detected significant COMP-Ang-1 protein levels only in the treated animal (**Figure 6E**). Full-length Tie-2 receptor levels were greatly reduced in the treated animal, a well-described downstream effect of Tie-2 activation.^23,31^ In order to control for standard protein loading as well as comparable proportions of endothelial cells, the endothelial marker eNos was used (**Figure 6E**). To confirm the immunoblot data, we performed immunoprecipitation (IP) using Fc-Tie2- captured COMP-Ang1, confirming COMP-Ang-1-expression (**Figure 6F**). Reciprocal IP for Tie2 demonstrated reduced Tie2-levels in the treated animals, consistent with receptor engagement, followed by ligand-induced internalization/degradation. Normal NHP lung tissue served as a control. We next investigated whether PDGFB levels could also serve as a secondary PD biomarker for COMP-Ang-1 induced Tie-2 activation in the treated NHPs. To assess PDGFB dynamics in NHP blood, we performed a Luminex assay after 4 hours of LNP002/mRNA-76 infusion. PDGFB levels were found to be reduced with all three doses (0.3, 0.6 and 0.9 mg/kg) at later time points (24 h, 48 h and 72 h) following the end of infusion (**Figure 6G**). Overall, in treated „healthy“ NHPs PDGF levels rather appear to decline over time in an almost dose-dependent manner, whereas VEGFA levels rather fluctuate over time and dose in contrast to steady levels observed in mice.(**Figure 6H**). These findings suggest investigating PDGFB as one potential blood-based PD biomarker reflecting lung specific Tie-2 activation under non-inflammatory (so called “target engagement”), healthy conditions for animals as a requirement for analyzing PK/PD in future dosing regimens in the context of GLP-toxicology studies to be conducted in rodents and NHPs, respectively.

**Figure 6.**
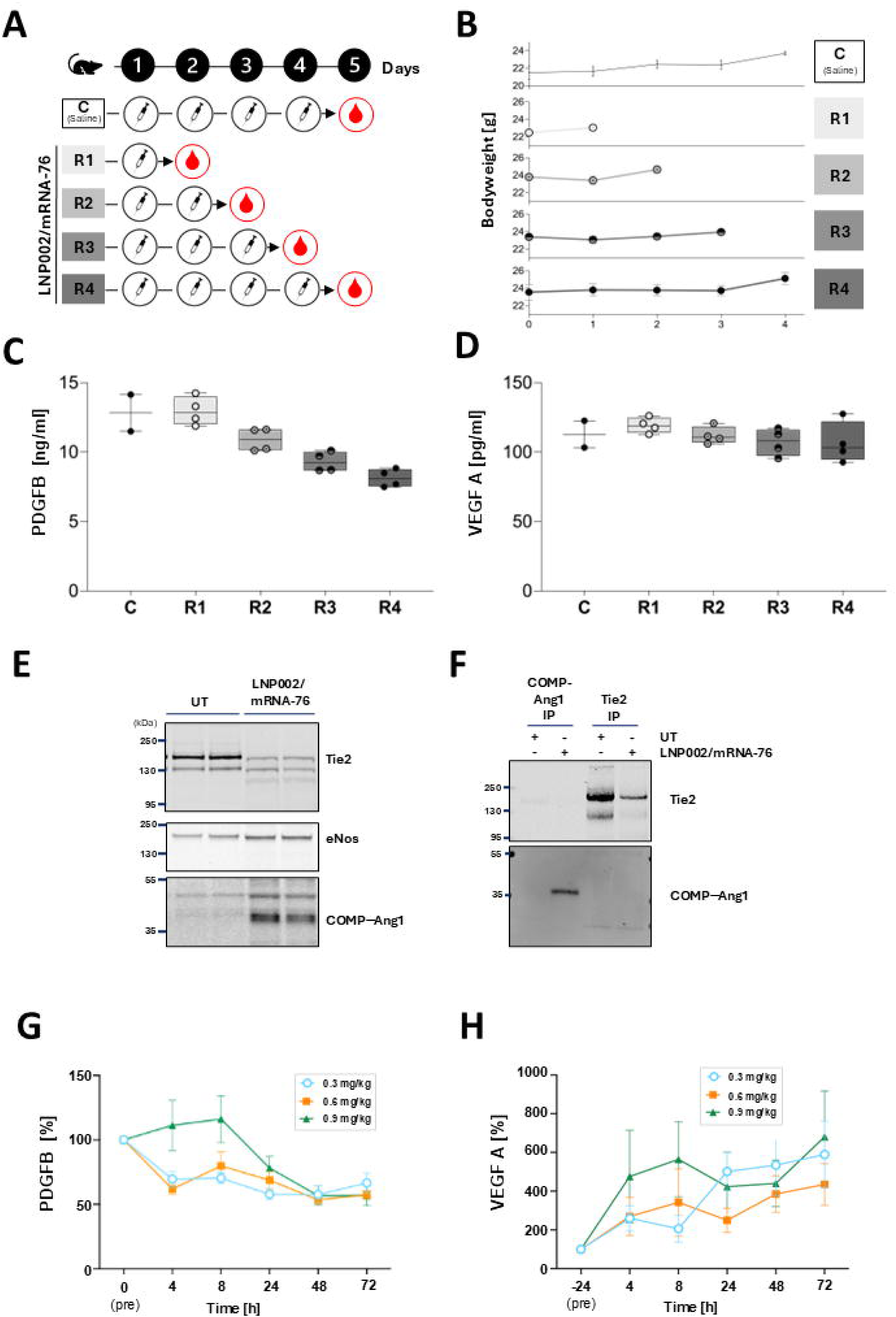
LNP002/mRNA-76 mediated expression of COMP-Ang-1 leads to secondary pharmaco-dynamic (PD) effects of Tie2 signaling in mice and in non- human primates (NHP) **(A)** Schematic of the repeated treatment schedule of mice (n=4) administered by tail vein injection (bolus) with LNP002/mRNA-76. Animals received LNP002/mRNA-76 at 0.75 mg/kg once (R1), twice (R2), three times (R3), or four times (R4). Blood samples were collected 24 h after the final administration. **(B)** Bodyweight of mice over the treatment period indicating a good tolerability of LNP002/mRNA-76. **(C), (D)** ELISA- based quantification of murine serum PDGFB levels and VEGF-A levels respectively following repeated dosing of LNP002/mRNA-76 in mice. **(E)** Immunoblot analysis of Tie2, eNos and COMP-Ang-1 antibodies on whole lung tissue lysates of an untreatedmale NHP in comparison to lung tissue obtained 24 h after a single 1 h infusion of LNP002/mRNA-76 (0.9 mg/kg). **(F)** Immunoblot analysis following immunoprecipitation of COMP-Ang1 and cynomolgus Tie2 from lung tissue obtained 24 h after a single 1 h infusion of LNP002/mRNA-76 (0.9 mg/kg). COMP-Ang1 was immunoprecipitated using Fc-Tie2, with a specific COMP-Ang1 signal detected exclusively in lung tissue in LNP002/mRNA-76-treated animals. Conversely, immunoprecipitation of Tie2 using Tie2- specific antibodies revealed reduced Tie2 levels in LNP002/mRNA-76-treated tissue compared with untreated control lung, consistent with ligand-induced Tie2 activation followed by receptor internalization and degradation. Commercially available normal cynomolgus lung tissue served as control. **(G), (H)** Luminex-based analysis of circulating PDGFB and VEGF levels in NHPs following a 4 h intravenous infusion of LNP002/mRNA-76 at the indicated dose levels. A time-dependent reduction of systemic PDGFB concentrations was observed at 0.3 and 0.6 mg/kg. Animals treated with 0.9 mg/kg exhibited a transient early increase in PDGFB followed by a decline to levels comparable to those measured at lower doses, supporting PDGFB as a COMP- Ang1/Tie2–dependent pharmacodynamic biomarker in vivo.

## Discussion

There are currently only four RNA/LNP-based drugs clinically approved. Following the 2018 approval of amyloidosis siRNA therapy Onpattro, an RSV vaccine (mRESVIA) and the two COVID-19 mRNA vaccines, Moderna (Spikevax) and Pfizer’s (Comirnaty) were approved in 2021 and 2024, respectively.^4,32–34^ All of these approved vaccines are based on neutral LNPs, containing ionizable lipids, hence producing a clinically validated platform and corroborating safety and efficacy for systemic nucleic acid delivery to the liver as well as for vaccination. This advance has subsequently enabled many therapeutic applications of small interfering RNA and messenger RNA modalities. Cationic lipids are effective at promoting cellular uptake and gene delivery, but cationic nanoparticles are also able to engage in interactions with cell membranes, resulting in membrane rupture and subsequent Ca2^+^ influx. This can potentially lead to cytotoxicity and cell death.^35,36^ This perception that cationic lipids are toxic led to the development of ionizable lipids with a neutral charge in the blood, as they were considered to be safer gene therapy delivery systems.^37,38^ However, there has been a recent increase in studies employing cationic lipid nanoparticles for delivery to the lungs.^39,40^

In the present study, we address critical translational concerns, including the occurrence of infusion-related immune reactions following systemic administration of positively charged LNPs formulated with permanently charged cationic lipids. To bridge this translational gap, we systematically evaluated how infusion duration affects immunological safety after systemic administration of a particular lung-targeting cationic LNP formulation, LNP002, in both mice and in NHPs. Historically, safety concerns related to cationic LNPs have primarily been reported in rodent studies involving bolus injections.^41–44^ By directly comparing short and prolonged infusion protocols in NHPs, we demonstrate that these responses are predominantly driven by peak-plasma exposure, rather than by intrinsic properties of cationic lipid chemistry. This contrasts with the well-documented and dose-limiting cytokine induction observed for nLNPs.^45,46^ In our data we show that there is no dose-dependent cytokine induction in NHPs following both the short and long infusion times of a cLNP. These data are consistent with previous observations showing that cytokine induction by positively charged, cationic LNPs is not the primary cause of dose-limiting toxicity in cationic LNP formulations.^47,48^ This mechanism appears to differ markedly from the adverse events associated with nLNPs, such as the approved Onpattro® (Patisiran, Alnylam Inc.), a systemically administered neutral LNP formulation delivering siRNA to hepatocytes, which requires extensive steroid premedication prior to administration to suppress cytokine-driven inflammatory reactions and other overactive immune responses.^49^

Our data indicates that under conditions of short infusion times, complement activation occurs in a dose-dependent manner and involves both the classical and the alternative complement activation pathways. Importantly, prolongation of infusion duration effectively mitigates complement activation, without altering total dose, nor compromising pharmacodynamic activity. This observation aligns with established animal models and clinical practice, where modulation of infusion kinetics is routinely employed to enhance tolerability and to mitigate infusion-related reactions.^50–52^ Particularly in efficacy mouse models, the commonly used bolus tail-vein administration has the disadvantage of generating high transient plasma concentrations of particle-based drug candidates, which can trigger adverse cytokine responses and fails to accurately mirror clinically relevant administration conditions. Beyond the non-physiologically relevant nature of bolus administration, our data suggests that mice are highly robust regarding complement and cytokine induction, making these murine observations poorly predictive of responses in other species. Hence, safety concerns for cLNPs and permanently positively charged lipids derived from rodent bolus studies should be considered with caution for the prediction of drug responses in other species, including humans. Our data therefore advocates for evaluating the safety of specific LNPs without bias, and case-by-case using relevant species, and clinically representative administration protocols.

These findings challenge the generally assumed poor intrinsic tolerability of cationic lipid formulations. From a broader perspective, our results suggest that the categorical exclusion of cationic LNPs from systemic RNA delivery strategies may be unwarranted, and even thwarting potential clinically useful candidates. Instead, careful consideration of exposure kinetics, particularly peak plasma concentration, should guide clinical translation. Prolonged infusion emerges as a practical and clinically feasible mitigation strategy that enables safe systemic administration while preserving the distinct lung-targeting properties of cationic nanoparticles.

To demonstrate the applicable clinical translatability of cLNPs we conducted a case study. The rationale for selecting mRNA-76, encoding for the Tie2-agonist COMP-Ang-1 as a drug candidate for a case study to illustrate the translational potential of LNP002 was fourfold. First, the organ tropism of cLNP002, especially its specific lung EC cell targeting, is consistent with the pathophysiology of early ARDS (acute respiratory distress syndrome) a life -threatening disease with high unmet medical need where fluid builds up on the lung. Secondly, ADRS is treated in the ICU setting. Unlike chronic indications, ARDS typically requires and enables only a short treatment window, necessitating one to three infusions per week, thereby allowing for controlled dosing and manageable exposure. Third, the Tie2 pathway, and its inactivation by Ang2, plays a well-validated role in promoting lung edema formation in patients with ARDS,^53^ providing a strong rationale for the innovative therapeutic strategy of mRNA-mediated expression of COMP-Ang-1 as the intended mechanism of action for this clinical approach (for more detail see also Radloff et al., 2023).^23^ Fourth, because of the technical challenges associated with producing potent Tie2-agonists, such as recombinant Ang-1,^26^ mRNA represents an attractive modality for the transient expression of Tie-2 agonists.

Establishing a robust, blood-detectable PD marker is a key prerequisite for clinical development of Tie2 agonist therapies. We originally hypothesized that secreted COMP-Ang-1 produced in endothelial cells would act through both autocrine and paracrine mechanisms, allowing its detection in blood samples as a primary pharmacodynamic marker. However, as noted previously, the very short half-life and predominant retention of COMP-Ang-1 within lung tissue prevented reliable quantification of the protein in serum or plasma from either mice or NHPs, despite clear evidence of local expression in lung tissue as detected by immunoblot in this study and in Radloff et al 2023.

In our previous studies we reported extensive preclinical pharmacodynamic data in mice for mRNA-76, which encodes the Tie2 agonist COMP-Ang-1. We employed the cationic LNP002/mRNA-76 formulation to demonstrate efficient and selective mRNA delivery to murine lung endothelial cells, enabling therapeutic modulation of endothelial signaling pathways involved in vascular barrier regulation and inflammatory lung injury.^23^

A limitation of our current study is that only GLP-like LNP002/mRNA-76 material was used in the NHP experiments, restricting the work to pharmacokinetic assessments. Because of this constraint, and the use of non-naïve NHPs, neither a formal toxicology evaluation, nor an extensive pharmacodynamic dataset could be generated at this stage. Despite the pharmacodynamic limitations in these NHP preclinical studies, we were able to demonstrate PD activity of LNP002/mRNA-76 for Tie2-signaling under non-inflammatory conditions by measuring the downregulation of PDFGB levels in blood samples of mice and NHPs. Repeated dosing of LNP002/mRNA-76 in mice resulted in a dose-dependent, systemic PDGFB reduction. Tie2-dependent PDGFB reduction was confirmed following a single 4 h LNP002/mRNA-76 infusion, consistent with Tie2 engagement and downstream signaling activity. This result is consistent with the expression pattern determined by murine scRNA-seq data to identify secreted proteins downstream of Tie2. ADAMTS1 and PDGFB were found to be selectively expressed in lung endothelial cells alongside the Tie2 receptor, which also represents the primary cell type to which mRNA-76 colocalizes. Other targets implicated in Tie2/FOXO-mediated transcriptional regulation, such as ADAMTS1, Endocan, CTGF, and Cyr61, identified from *in vitro* tissue culture experiments using cells derived from patients carrying Tie2-activating mutations^54–57^ could not be validated as PD markers in our *in vivo* experiments. These findings suggest that PDGFB may serve as a viable, blood-based PD biomarker for Tie2-modulating drugs particularly targeting the lung vasculature, likely due to its relatively restricted expression in this tissue. However, establishing a robust, blood-detectable PD marker remains a key prerequisite with respect to determination of therapeutic dosing for future clinical development of Tie2 agonist therapies.

Building directly on the previously established pulmonary endothelial targeting and efficacy of LNP002/mRNA-76,^23^ we evaluated whether modulation of infusion kinetics could reduce immune-response-related complement activation, while maintaining pharmacodynamic activity. Our findings provide mechanistic insights into the nature of infusion-associated reactions with cLNPs and identify prolonged infusion as an effective and clinically feasible strategy to enable their safe systemic administration in NHPs in the absence of immunosuppressive premedication, thereby supporting their translational potential for pulmonary endothelial-targeted RNA therapeutics.

## Materials & Methods mRNA synthesis

Codon optimized, CleanCap-formulated mRNAs were synthesized and purified by BioSpring GmbH (Frankfurt am Main, Germany) and AmpTec (now Merck, Hamburg, Germany). These mRNAs contained the 5′ UTR and signal peptide sequence of human MCP1 (NM_002982.3), the 3′ UTR sequence of human vWF (NM_000552.4), and the coding sequences for COMP Ang1 (mRNA 76), Firefly luciferase (mRNAFLUC), and enhanced green fluorescent protein (mRNAeGFP). In all constructs, uridine was globally substituted with N1 methylpseudouridine.

## LNP preparation and characterization

The cationic lipid (L-arginyl)-L-2,3-diaminopropionic acid-N-palmityl-N-oleyl-amide (Pantherna Therapeutics, Hennigsdorf, Germany), the helper lipid 1,2-diphytanoyl-sn-glycero-3- phosphoethanolamine (Corden Pharma, Plankstadt, Germany), and the PEGylated lipid 1,2- distearoyl-sn-glycero-3-phosphoethanolamine-N-[methoxy(polyethylene glycol)-2000] (sodium salt) (Corden Pharma, Plankstadt, Germany) were dissolved in absolute ethanol at a molar ratio of 50:49:1. The ethanolic lipid phase was combined with mRNA dissolved in 10 mM sodium citrate buffer (pH 5.5), yielding a lipid-to-mRNA mass ratio of 20:1. The final mRNA concentration in the formulations ranged from 100 to 280 µg/ml. For nanoparticle formation, aqueous mRNA solutions and lipid solutions in ethanol were mixed at a volume ratio of 2:1 (aqueous:ethanol) using a microfluidic mixer (NanoAssemblr®, Precision Nanosystems, Vancouver, Canada) operated at a total flow rate of 18 ml/min. Following particle formation, the formulations were dialyzed for a minimum of 18 h at 4 °C against 10 mM Tris buffer (pH 7.4) containing 9 % sucrose using Slide-A-Lyzer dialysis cassettes (MWCO 3.5 kDa, Thermo Scientific, Rockford, IL, USA). Particle size distribution and zeta potential were determined by dynamic light scattering (Zetasizer Nano ZS, Malvern Instruments, Malvern, UK). RNA encapsulation efficiency and total RNA content were quantified using the Quant-iT RiboGreen RNA Assay Kit (Thermo Fisher Scientific, Waltham, USA). The resulting lipid nanoparticles exhibited a Z-average diameter of 80–90 nm, a polydispersity index (PDI) below 0.1, and a zeta potential ranging from +5 to +10 mV when suspended in 10 mM Tris buffer (pH 7.4) supplemented with 9% sucrose. Encapsulation efficiency consistently exceeded 95 %. All formulations were stored at –80 °C until further use.

### Cell transfection *in vitro*

HeLa cells were cultivated in T75 culture flasks at 37 °C, 5 % C02 in DMEM Glutamax (Gibco, cat. #61965-026) supplemented with 10 % FCS and 1 % P/S. Cells were washed with PBS and after a 3 min incubation with 3 ml 0.25 % trypsin-EDTA (Gibco, cat. #25200072) for 3 min cells were detached from the flask by swiveling and tapping. Trypsinization was stopped using 3 ml supplemented cell culture medium. Cell suspension was transferred into a 15 ml tube and spun down. The cell pellet was resuspended in fresh culture medium before cell counting via Countess III Cell Counter with Trypan Blue. For passaging 10 % of the cell suspension was transferred into a fresh 75 cm^2^ culture flask (Thermo Scientific, cat. #156499) and for in vitro screens 100.000 cells per well were plated in 12-well cell culture plates (Thermo Scientific, cat. #140675) containing 1 ml culture medium. Cells were transfected after 24-48 h when cells reached a confluency of 60-70 %. LNP were thawed at RT in a water bath and homogenized by flicking the tube. LNP amounts of (0.25, 0.5 and 1.0 µg) for cLNPs and (2, 10, 25 µg) for nLNPs per well were added in a dropwise fashion and the plate was gently swirled to disperse the LNPs within the well. After a 24 h incubation fluorescent images were acquired with a Nikon Eclipse Ti-U equipped with an ELWD S Pan Fluor 40×/0.6 objective lens.

## *In vivo* biodistribution study and repeated dosing study in mice

Mice experiments were conducted at EPO (Experimental Pharmacology & Oncology Berlin-Buch GmbH) in accordance with the German Animal Protection Law and the EU guideline European Convention for the Protection of Vertebrate Animals Used for Experimental and Other Scientific Purposes (ETS 123), as well as the German Animal Protection Law – Version July 2014 (Tierschutzgesetz: zuletzt geändert durch Art. 3 G v. 28.7.2014 I 1308) and the Regulation on the Protection of Animals Used for Experimental or Other Scientific Purposes (Tierschutz-Versuchstierverordnung, TierSchVersV: geändert durch Art. 6 V v. 12.12.2013 I 4145).

Compliance was monitored by the local authorities (Landesamt für Gesundheit und Soziales, LAGeSo). Male C57Bl/6NRj mice (SOPF), aged 8–10 weeks, were obtained from Janvier Labs (France) and housed under standard laboratory conditions with a 12 h/12 h light–dark cycle. Animals were kept in groups of three with ad libitum access to tap water and a standard rodent diet. LNPs at a concentration of 300 µg/mL were stored at −80 °C and thawed in a water bath to room temperature prior to use. All injections were performed at room temperature. For the biodistribution study, three mice per group received an intravenous (i.v.) bolus injection of 125 µL LNP per 25 g body weight via the tail vein, corresponding to an mRNA dose of 1.5 mg/kg. Control animals received an equal volume of isotonic 0.9 % NaCl solution. Six hours after injection, mice were euthanized and lung and liver tissues were dissected and snap-frozen. For the repeated-dosing study, four mice per group received 0.75 mg/kg LNP002/mRNA-76 once every 24 hours, following the study design shown in Figure 6A. Bodyweight was monitored daily. Mice were euthanized 16 hours after the final dose, and serum samples were collected and snap-frozen for later analysis.

## Luciferase activity assay in murine lung and liver tissue

The biodistribution and expression efficiency of various LNP formulations containing firefly luciferase (Fluc)–encoding mRNA were assessed ex vivo by quantifying relative luciferase activity using the Luciferase Assay System (Promega, cat. #E1500). Tissues were cut into small pieces, weighed (20–100 mg), and lysed in chilled 1× Cell Culture Lysis Reagent (Promega, cat. #E1531) at a ratio of 10 µL lysis reagent per mg tissue. Following bead mill homogenization, lysates were centrifuged, and 20 µL of the protein containing supernatant was combined with 100 µL of assay reagent for immediate luminescence measurement. Relative luminescence units (RLU) were normalized to total protein content, determined using the Pierce™ BCA Protein Assay (Thermo Scientific, cat. #23227).

## Western Blot Assay

Tissue lysates mixed with NUPAGE sample buffer were heated at 70 °C for 10 min and loaded onto NuPAGE 4–12 % Novex Bis-Tris Gels for SDS-PAGE electrophoresis in MOPS buffer (all from Invitrogen, cat. #NP0335BOX, NP0001). Proteins were transferred to nitrocellulose membrane (Cytiva, cat. #10600003; NuPAGE™ Transfer Buffer (Invitrogen, cat. #NP00061). After a 60 min blocking step (Intercept® Blocking Buffer; LiCor, cat. #927-60001) membranes were probed at 4 °C overnight with specific primary antibodies diluted in Intercept Antibody Diluent (LiCor, cat. #927-65001) followed by washing step with Tris Buffered Saline with Tween® 20 (CST, cat. #9997S). Binding of primary antibodies was visualized using near infrared IRDye secondary antibodies (LiCor; cat. #926 32213, #926 32214, #926 68072), and fluorescence signals were acquired with the LiCor Odyssey CLx system. Band intensities were quantified using Empiria Studio Software v2.1.0134 (LiCor). The primary antibodies used included Angiopoietin 1 (Abcam, cat. #ab183701), human Tie2 (Cell Signaling Technology, cat. #7403), mouse Tie2 (Merck, cat. #05 584), and eNOS (Cell Signaling Technology, cat. #32027).

## Immunoprecipitation

Frozen control cynomolgus lung tissue (Hölzel Biotech, cat. # CLB-NHP-TI117) and lung tissue of an mRNA-76 treated animal (provided by Charles River) was weight and cut into fragments of 30-100 mg and homogenized in 20 µl/mg chilled Pierce™ IP lysis buffer (Thermo Scientific, cat. #87788) via TissueLyser LT bead mill (Qiagen, cat. #69980). After centrifugation protein content of the supernatants was measured using Pierce™ BCA kit. Lysate volume was adjusted to ensure equal total protein content in all samples. Immunoprecipitation was performed using Invitrogen’s Antibody Dynabead Coupling Kit (Thermo Scientific, cat. #14311D) according to manufactureŕs instructions. In brief, Anti-Tie2 antibody ab33 (R&D Systems, Merck, cat. #05- 584) for Tie2 IP or Tie2-Fc fusion protein (R&D Systems, cat. #313-TI) for COMP-Ang1 IP were covalently coupled to M-270 Epoxy Dynabeads™ (Thermo Scientific, cat. #14311D) and 1.5 mg of antibody-coupled beads were incubated with tissue lysates overnight at 4 °C. Beads were washed with ice-cold Pierce™ IP lysis buffer, after denaturation with 1x NUPAGE LDS-sample buffer containing 50 mM Dithiothreitol and Halt™ Protease and Phosphatase Inhibitor Cocktail (all Thermo Scientific, cat. #NP0007, #NP0009, #78441) Dynabeads were removed by centrifugation. Immunoprecipitated lysates and input lysates were analysed by Western blotting analysis.

## Single cell RNA sequencing

Single cell suspensions generated from fresh mouse lung samples were loaded onto the chip G (10x Genomics) and processed according to the Chromium Next GEM Single Cell 3’ workflow using the Chromium Controller device (10x Genomics). For each sample, 11000 single cells were loaded, with an average cell viability of 88 %. The resulting libraries were sequenced using a NextSeq2000 device (Illumina). For the analysis of the sequencing data a new reference was constructed based on the GRCm39 genome and the GENCODE M29 annotation which included the mRNA-76 transgene contigs along with their annotation and was then filtered according to the standard recommendations of 10x CellRanger (v7.0.0) and packaged for use by the ‘cellranger mkref’ command using default parameters. All sequencing runs were demultiplexed with ‘cellranger mkfastq’ using default parameters. The demultiplexed samples were processed with ‘cellranger count’ using the non-default parameter ‘--expect-cells=5000’.^58,59^

## NHP animal studies

Two independent non-GLP pharmacokinetic studies were conducted in male Cynomolgus monkeys to evaluate the impact of infusion duration on systemic mRNA exposure and infusion- related immune activation. In the 1 h-Infusion study (Charles River Laboratories; Study No. 20245723), cationic LNP formulation (LNP002) encapsulating mRNA-76 were administered as a single 1 h intravenous infusion. Animals (n = 2 per group) received 0.3, 0.6 or 0.9 mg/kg. An untreated control group was included. The infusion volume was 10 mL/kg. In the 4 h-Infusion study (Labcorp; Study No. 8473-792), LNP002/mRNA-76 was administered as a single 4 h intravenous infusion at 0.3, 0.6 or 0.9 mg/kg (n = 5 males per dose group). Infusions were delivered via calibrated external pumps with a target dose volume of 15 mL/kg. The nominal mRNA dose levels were the same to those used in the 1 h infusion study, enabling direct cross- study comparison. All procedures complied with institutional animal welfare regulations and study-specific SOPs. Blood samples were collected pre-dose and at predefined time points up to 48 h (1 h study) or up to 72 h (4 h study) post-infusion. Plasma (4 h study) or serum (1 h study) concentrations of PAN004 mRNA were quantified using an RT-qPCR–based assay specific for mRNA-76. Pharmacokinetic parameters were derived using compartmental modeling (PKSolver v2.0 in the 1 h study). Complement activation was assessed by quantification of C3a, Bb (alternative pathway), and soluble C5b-9 (terminal complement complex) at predefined post- infusion time points. Systemic cytokine and chemokine responses were evaluated using multiplex immunoassays, including IL-1β, IL-2, IL-4, IL-5, IL-6, IL-8, IL-10, IL-17A, IFN-γ, TNF-α, and MCP-1.

Pharmacokinetic parameters, complement markers, and cytokine concentrations were summarized descriptively. Individual concentration–time profiles were evaluated for infusion duration–dependent differences in C_max_, time to peak concentration (T_max_), area under the curve (AUC), and terminal half-life. Comparisons between 1 h and 4 h infusion cohorts were performed on a descriptive basis due to differences in group size and study conduct (n = 2 vs. n = 5 per dose group). For dose-dependent trends within individual studies, fold changes relative to pre-dose values were calculated for complement and cytokine markers. No formal hypothesis testing was prespecified. Graphical analyses and concentration–time visualizations were performed using standard scientific plotting software.

## Data availability statement

The authors confirm that the data supporting the findings of this study are available from the corresponding author upon reasonable request. The scRNA-Seq data sets are deposited at GEO Datasets: GSE241514.

## Acknowledgements

This work was funded by Pantherna Therapeutics GmbH. We want to thank Professor D. Riesner for his generosity and support. We acknowledge Volker Fehring and Akansha Moga for their formulation support. We also acknowledge the work of Katharina Ahrens in proofreading this paper. Nano Banana 2 was used in part for generation of the graphical abstract.

