## Supplemental Figure for "Translational Pharmacokinetics and Pharmacodynamics of a Cationic mRNA–Lipid Nanoparticle from Mice to Non Human Primates"

### Supplement

A

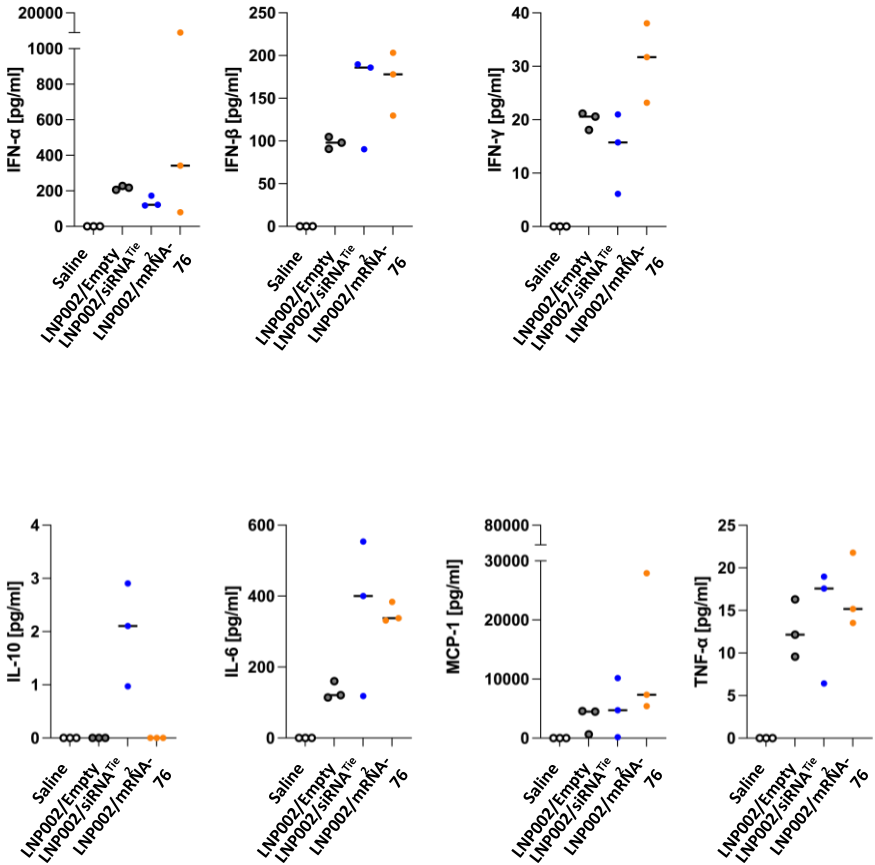

B

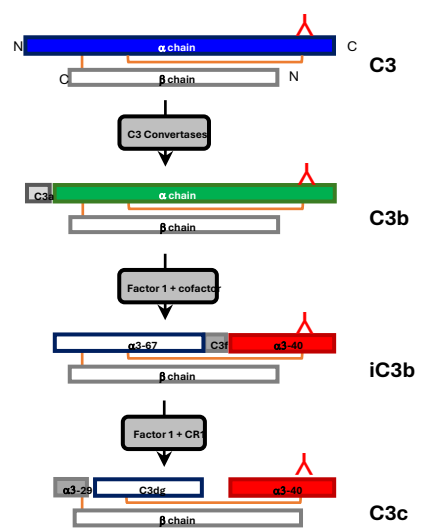

C

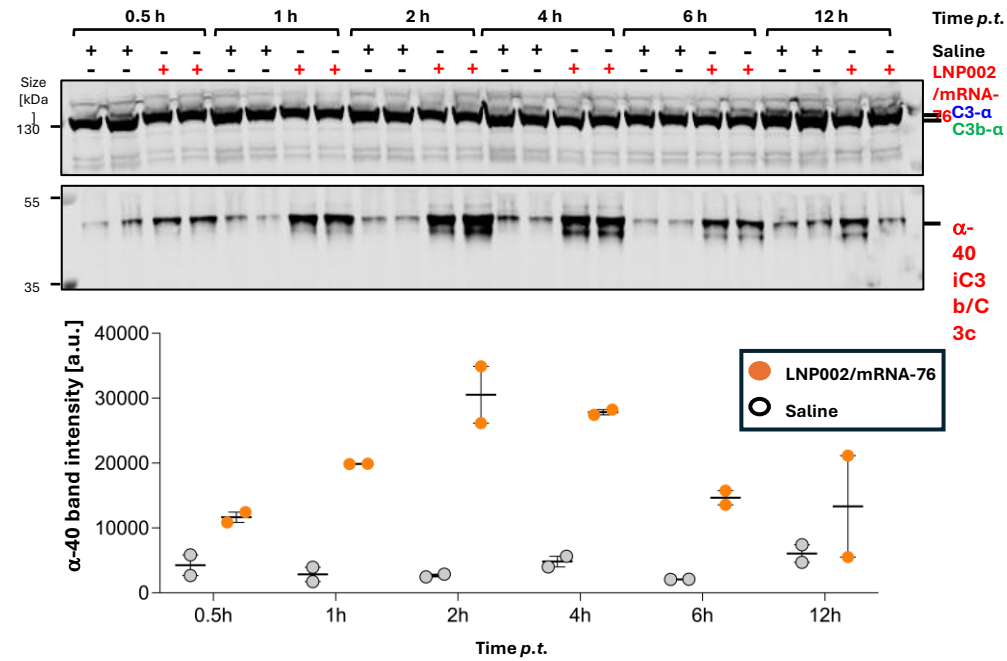

**Supplemental Figure 1. Cytokine response and complement activation in mice after i.v. bolus tail vein administration of LNP002/mRNA-76**

**(A)** Serum cytokine level of mice 6 h post bolus treatment with saline, LNP002/empty (LNP002 without a nucleic acid cargo), LNP002/siRNA<sup>Tie2</sup> and LNP002/mRNA-76. Animals (n=3) received doses of nucleic acid mRNA-76 at 2mg/kg, siRNA at 2mg/kg or with corresponding lipid mixtures for empty LNPs and saline volume as controls **(B)** Schematic drawing of activation and inactivation of C3 complement. The different C3 complement cleavage products detected by the C-terminal specific antibody Anti-C3 antibody (EPR19394; abcam) are indicated. **(C)** Immunoblot with Anti-C3 antibody on serum from mice collected at different time points post treatment (n=2). Mice were treated by bolus injection with saline or LNP002/mRNA-76 (2mg/kg). The C3 alpha change and the corresponding cleavage products recognized by the C-terminal specific antibodies are indicated by arrows.
